# Essentiality, Protein-Protein Interactions and Evolutionary Properties are Key Predictors for Identifying Cancer Genes Using Machine Learning

**DOI:** 10.1101/2021.09.01.458494

**Authors:** Amro Safadi, Simon C. Lovell, Andrew J. Doig

**Affiliations:** Division of Evolution and Genomic Sciences, School of Biological Sciences, Faculty of Biology, Medicine and Health, The University of Manchester, Manchester M13 9PT, United Kingdom; Division of Neuroscience and Experimental Psychology, School of Biological Sciences, Faculty of Biology, Medicine and Health, The University of Manchester, Manchester M13 9PT, United Kingdom

## Abstract

The identification of genes that may be linked to cancer is of great importance for the discovery of new drug targets. The rate at which cancer genes are being found experimentally is slow, however, due to the complexity of the identification and confirmation process, giving a narrow range of therapeutic targets to investigate and develop. One solution to this problem is to use predictive analysis techniques that can accurately identify cancer gene candidates in a timely fashion. Furthermore, the effort in identifying characteristics that are linked to cancer genes is crucial to further our understanding of this disease. These characteristics can be employed in recognising therapeutic drug targets. Here, we investigated whether certain genes’ properties can indicate the likelihood of it to be involved in the initiation or progression of cancer. We found that for cancer, the essentiality scores tend to be higher for cancer genes than for all protein coding human genes. A machine-learning model was developed and we found that essentiality related properties and properties arising from protein-protein interaction networks or evolution are particularly effective in predicting cancer-associated genes. We were also able to identify potential drug targets that have not been previously linked with cancer, but have the characteristics of cancer-related genes.

**Author Summary:** Mutations in numerous genes are known to be involved in cancer, yet there are undoubtedly many more to be discovered. We analysed a set of hundreds of cancer genes with the aim of finding out what makes them different from genes not known to be mutated in cancer. In particular, we found that genes that are essential for the survival of an organism are more likely to be involved in cancer. We used the gene properties that we examined to develop an artificial intelligence method that can accurately predict whether a gene is involved in cancer or not. Applying the method gives hundreds of non-cancer genes that resemble cancer genes. New discoveries of cancer genes are likely to be found within this set.

## Introduction

The identification of cancer-related genes (referring to both oncogenes and tumour suppressor genes) remains a key challenge. Among all human genes, approximately 3.5% have been directly implicated in cancer initiation and progression [1], although it is likely that many more remain to be found. Accurate identification of genes potentially related to cancer would provide an opportunity to advance both personalised treatment of cancer and aid drug discovery by providing new targets. The Cancer Gene Census of COSMIC [1] provides an expert-curated dataset of cancer-associated genes, relying on tumour sample analysis to identify cancer genes. However, the expert-curation is a lengthy and complex process due to several factors including the availability of tumour samples and the difficulty in sequencing them. Several studies have previously focused on developing algorithms to identify cancer genes and their mutations. Most studies are based on the frequency or pattern of mutations in multiple tumours with some analysing the characteristics of these genes [2, 3]. The goal of accurately predicting cancer genes still eludes us.

One viable approach may be to enrich the set of properties that characterise these genes and combine their properties to reach a more reliable prediction method. A number of characteristics may be correlated with the likelihood of a gene being associated with cancer. A prime candidate is essentiality. A gene is considered essential when loss of its function compromises the viability of an individual [4]. Essentiality is a quantitative measure and not a simple divide between essential versus non-essential, as defining it as such would be impossible due to the changeable nature of essentiality based on the genetic and environmental context. The identification of essential genes in multiple organisms has provided researchers with vital insights into the mechanisms of biological processes [5]. For example, essential genes are likely to encode hub proteins in protein–protein interaction networks, signifying more interacting partners than non-essential genes. Furthermore, essential genes are more likely to be abundantly and ubiquitously expressed in cells and tissues and have smaller-sized introns [5]. Several studies determined the relationship between evolutionarily conservation and the degree of essentiality in genes with variations in findings between species [6, 7]. Findings from human genes point to an association where the more essential the gene is, the less likely it is to show enrichment of missense mutations. In contrast, the number of synonymous mutations is not dependent on essentiality. This indicates that purifying selection acts more stringently on essential genes [4].

One could argue that genes implicated in driving and initiating tumours, which generally do not compromise viability in a direct manner, are thus unlikely to score high on the essentiality spectrum. However, there are indications that human genes associated with genetic disease are likely to be essential [5]. Cassa et al [8] investigated heterozygous protein-truncating variants in over 60,000 individuals from the Exome Aggregation Consortium (ExAC) dataset [9] using the ‘shet’ essentiality score (a metric that provides Bayesian estimates of the selection coefficient against heterozygous loss-of-function variation) and were able to predict phenotypic severity, age of onset and penetrance for Mendelian disease-associated genes. In addition, genes involved in neurological phenotypes, including autism, congenital heart disease and inherited cancer risk, seem to be under more intense purifying selection, which may indicate essentiality. Quantitative estimates of essentiality thus appear to be useful in Mendelian disease gene discovery efforts.

Here, we identify combinations of gene properties that have not been previously used to assess the likelihood of a gene to be cancer-associated. We study whether cancer-associated genes are more likely to be essential than non-cancer genes and check whether an uplift in predicting a gene to be a cancerous can be achieved by using essentiality-related properties. These findings might also indicate if these genes are more likely to be under stronger selection than other non-cancer related genes. We were able to build an accurate machine-learning model to predict cancer genes, where essentiality-related properties were particularly useful. Using this machine learning approach, we were able to identify further candidate genes for cancer, in addition to those currently reported in COSMIC census (October 2018).

## Results

### Cancer Genes and Essentiality Scores

We first determined whether cancer-related genes are likely to have high essentiality scores. We aggregated several essentiality scores calculated by multiple metrics [4] for the list of genes identified in the COSMIC Census database (Oct 2018) and for all other human protein coding genes. Two different approaches to scoring genes’ essentiality were reviewed and compared in [4]. The first group of methods calculates the essentiality scores by measuring the degree of loss of function caused by a change (represented by variation detection) in the gene. It uses the following methods: residual variation intolerance score (RVIS), LoFtool, Missense-Z, the probability of loss-of-function intolerance (pLI) and the probability of haplo-insufficiency (Phi) (S1 Table). The second group (Wang, Blomen and Hart-EvoTol) studies the impact of variation on cell viability. For all methods above measuring essentiality, a higher score indicates a higher degree of essentiality, with the exception of Missense-Z21 where a lower score indicates higher degree of essentiality.

We find that on average the cancer genes exhibit a higher degree of essentiality compared to the average scores calculated for all protein coding human genes and all metrics. We find that genes associated with cancer have higher essentiality scores on average in both categories (intolerance to variants and cell line viability) compared to the average scores across all human genes (data can be found in S2 Table), with p-values consistently < 0.00001 (Table 1). We also investigated whether Tumour Suppressor (TS) genes as a distinct group of genes would show different degrees of essentiality. We found that no significant difference in the degree of essentiality on average for that group compared to the set of all cancer genes (Table 1).

**Table 1.** Essentiality scores comparisons between cancer and all human genes. All p-values that compare cancer genes or TS genes to all genes are <0.00001.

| Method | Mean<br>Essentiality | Mean<br>Essentiality | Ratio<br>(cancer | Mean<br>Essentiality | Ratio (TS<br>genes/all<br>genes) |
| --- | --- | --- | --- | --- | --- |

|  | Score for all<br>genes | Score for cancer<br>genes | genes/all<br>genes) | Score for TS<br>genes |  |
| --- | --- | --- | --- | --- | --- |
| Phi | 0.27 | 0.63 | 2.3 | 0.58 | 2.2 |
| Wang | 0.42 | 0.62 | 1.5 | 0.65 | 1.6 |
| S_het | 0.06 | 0.12 | 2.1 | 0.12 | 2.1 |
| LofTool | 0.50 | 0.70 | 1.4 | 0.71 | 1.4 |
| Missense-Z | 0.69 | 1.86 | 2.7 | 1.85 | 2.7 |
| RVSI | 50.0 | 68.3 | 1.4 | 68.9 | 1.4 |

The results are particularly of interest in the context of cancer, as essential genes have been shown to evolve more slowly than nonessential genes [6, 10, 11] although some conflicts have been reported [11]. A slower evolutionary rate indicates less probability to evolve resistance to a cancer drug. This is particularly important in the case of anticancer drugs as it was reported that these drugs cause a change in the selection pressure when administrated, leading to increased drug resistance [12].

### Modelling Results

This association between cancer-related genes and essentiality scores prompted us to develop methods to identify cancer-related genes using this information. We used a machine-learning approach, using the DataRobot platform. This platform applies a range of open-source algorithms and optimizes the weights of terms to produce the most accurate classifier. We focused on properties related to protein-protein interaction networks, as essential genes are likely to encode hub proteins, i.e. those central to the network [10, 13].

The model development workflow steps (i.e., the model blueprint) is shown in Fig 1:

**Fig 1.**
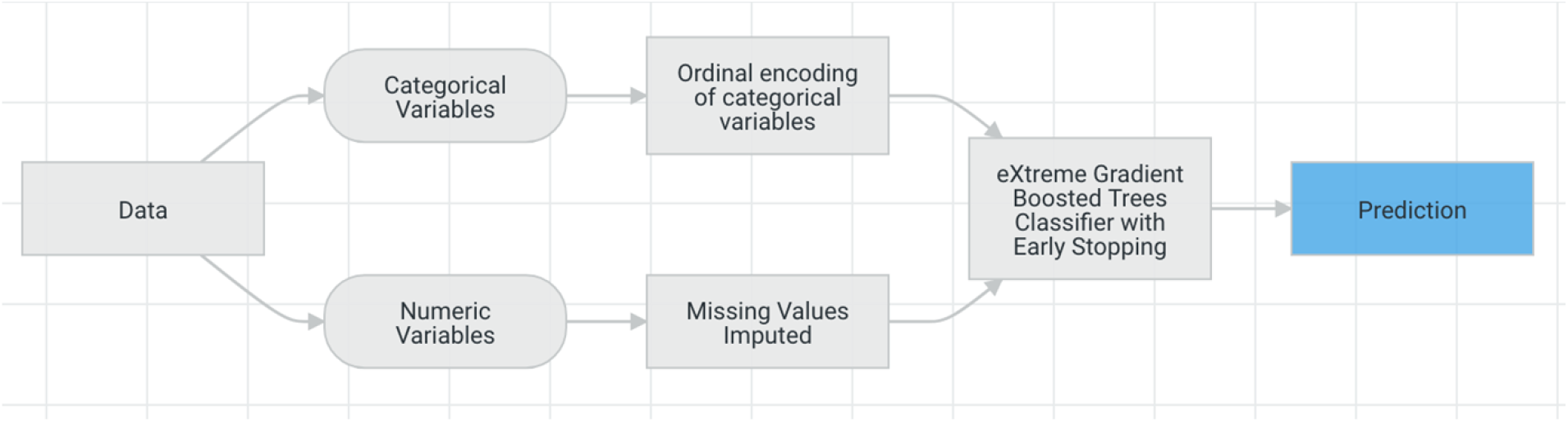
Modeling stages.

The model blueprint (Fig 1) is the result of trying multiple configurations and selecting the one that yields the best performance. It shows the preprocessing steps and the algorithm used in our final model and illustrates the steps involved in transforming input into a model. In this diagram, ‘Ordinal encoding of categorical variables’ converts categorical variables to an ordinal scale while the ‘Missing Values Imputed’ node imputes missing values. Numeric variables with missed values were imputed with an arbitrary value (default -9999). This is effective for tree-based models, as they can learn a split between the arbitrary value (−9999) and the rest of the data (which is far away from this value). A log showing any data transformation and imputation of values can be found in S4 Table.

A total of ten different modeling approaches (or blueprints) were run on the data to ensure the selection of the best performing approach (the list of these can be found in S3 Table along with their performance metrics). The performance metric used to rank the models was Logarithmic Loss (LogLoss), LogLoss is an appropriate and known performance measure when the model is of a binary-classification type. The LogLoss measures confidence of the prediction and estimates how to penalise incorrect classifications. The selection mechanism for the performance metric takes the type of model (binary classification in this case) and distribution of values into consideration when recommending the performance metric. However, other performance metrics were also calculated and can be found in the S3 Table. The performance metrics are calculated for all validation sets to ensure that the model is not over-fitting (Table 2). The model with the best performance result was eXtreme Gradient Boosted Trees Classifier with Early Stopping. This model was developed in a project created with version 7.0 of DataRobot platform. The model shows very close LogLoss values for training/validation and holdout (20% of the data was left out of the model training and validation datasets to be used as a blind test) data sets, ensuring no over-fitting.

**Table 2.**
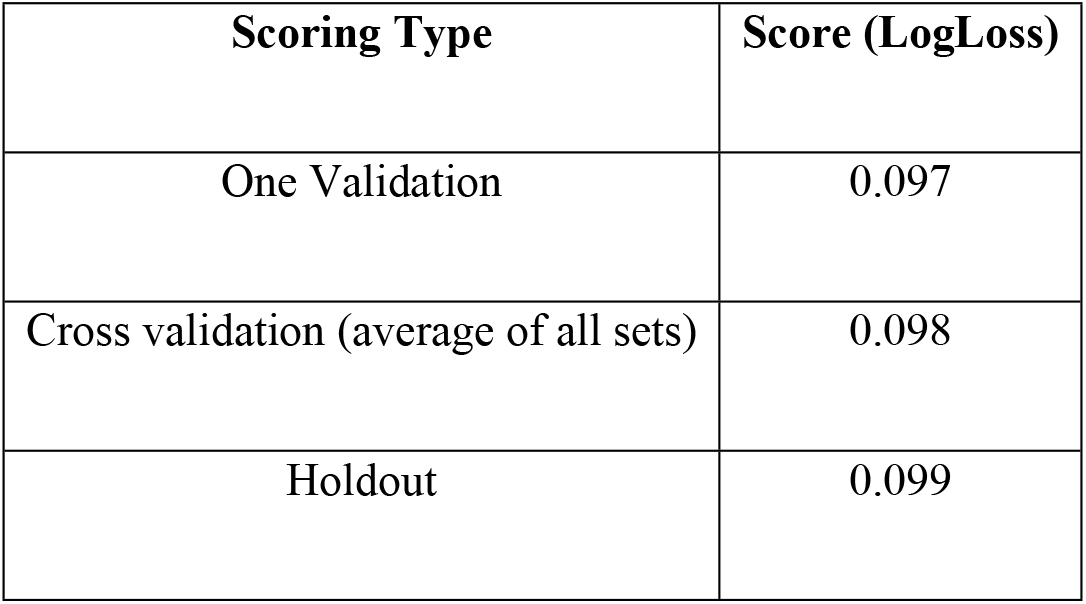
Performance metric (LogLoss) scores for our model validation and holdout segments. Cross Validation is the average of all 5 scores for each validation set.

To demonstrate the effectiveness of our model, a chart was constructed (Fig 2) that shows across the entire validation dataset (divided into 10 segments or bins and ordered by the average outcome prediction value), the average actual outcome (whether gene has been identified as cancer gene or not) and the average predicted outcome for each segment of the data (ordered from lowest average to highest per segment). By showing the actual outcomes alongside the predictive values for the dataset, we can see how close these predictions are to the actual known outcome for each segment of the dataset. We can also determine whether the accuracy diverges in cases where the outcome is confirmed cancer or when it is not, as the segments are ordered by their average of outcome scores. In general, the steeper the Actual line is, and the more closely the Predicted line matches the actual line, the better the model. A close match between these two lines is indicative of high predictive accuracy; a consistently increasing line is another good indicator of satisfactory model performance. The graph we have for our model indicates that our prediction model is highly accurate.

**Fig 2.**
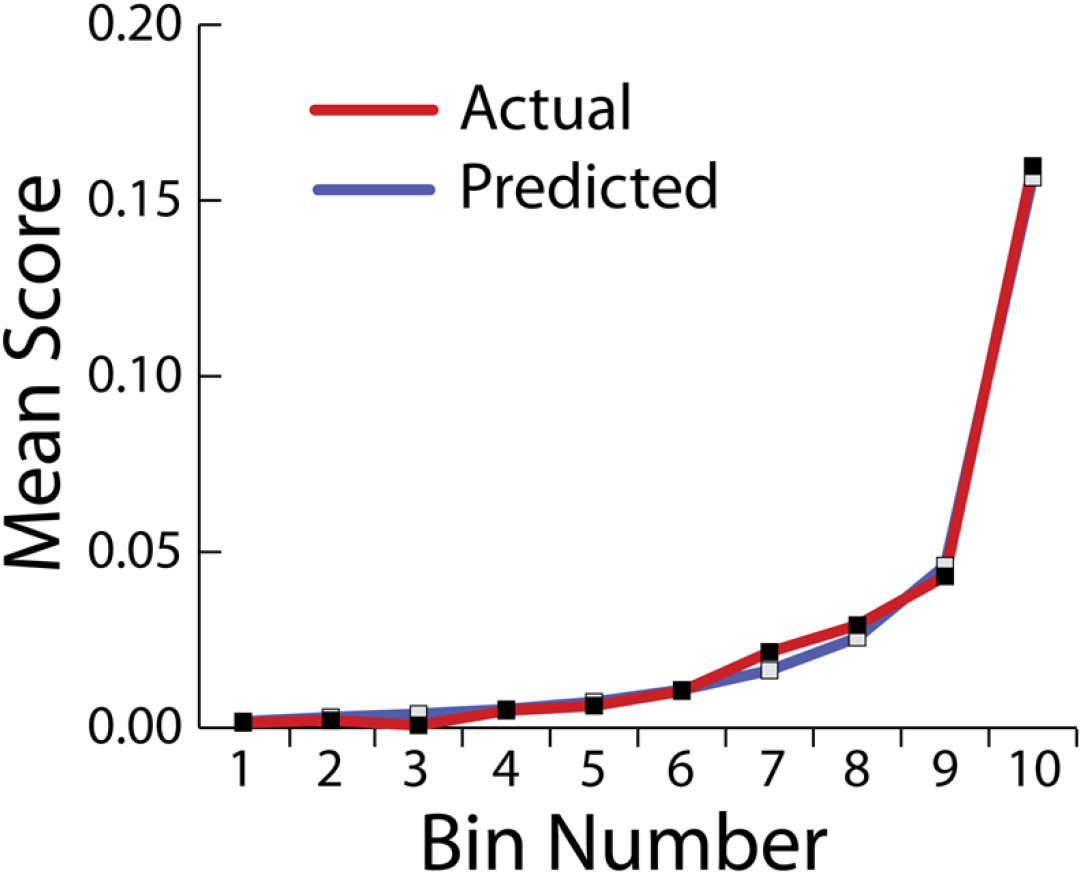
Lift Chart illustrating model accuracy. The left side of the curve indicates where the model predicted a low score for likelihood of being a cancer gene, while the right side of the curve indicates where the model predicted a high score. The “Predicted” blue line displays the average prediction score for the rows in that bin. The red “Actual” line displays the actual percentage for the rows in that bin.

The confusion matrix (Table 3) and the summary statistics (Table 4) show the actual versus predicted values for both true/false categories for our training dataset (80% of the total dataset). The model statistics show the model reached just over 89% specificity and 60% sensitivity in predicting cancer genes. This means that we are able to detect over half of cancer genes successfully while only misclassifying around 10% of non-cancer genes within the training/validation datasets. Both the F1 score (harmonic mean of the precision and recall) and Matthews Correlation Coefficient (MCC is the geometric mean of the regression coefficient) indicate high level of accuracy.

**Table 3:** Confusion Matrix. True Positives (TP); True Negatives (TN); False Positives (FP); False Negatives (FN), where Positives are cancer genes and Negatives are non-cancer genes.

|  |  | Predicted |  |  |
| --- | --- | --- | --- | --- |
|  |  | - | + |  |
| Actual | - | 12493 (TN) | 1490 (FP) | 13983 |
|  | + | 159 (FN) | 243 (TP) | 402 |
|  |  | 12652 | 1733 |  |

**Table 4.** Summary statistics for model performance.

| F1 Score | True Positive Rate (Sensitivity) | False Positive Rate (Fallout) | True Negative Rate (Specificity) | Positive Predictive Value (Precision) | Negative Predictive Value | Accuracy | Matthews Correlation Coefficient |
| --- | --- | --- | --- | --- | --- | --- | --- |
| 0.23 | 0.61 | 0.11 | 0.89 | 0.14 | 0.99 | 0.89 | 0.25 |

### The False Positives

The 1490 false positives are genes that the model classifies as cancer genes, even though they are not. Newly discovered cancer genes are likely to come from this set. In order to further confirm the model’s ability to predict cancer genes, we used it on 190 cancer genes that had been added to COSMIC’ Cancer Census Genes between October 2018 and April 2020. We were able to predict 56 genes out of the newly added 190 genes as cancer genes, all of which were among the false positives detected by the model. This indicates that the model is indeed suitable to use to predict novel candidate cancer genes that could be experimentally confirmed later. Specificity and sensitivity levels were similar to the earlier set.

Another way to visualise the model performance, and determine the optimal score to use as a threshold between cancer and non-cancer genes, is the Prediction Distribution graph (Fig 3) that illustrates the distribution of outcomes. The distribution (in purple) shows the outcome where gene is not classified as a cancer gene while the second distribution (in green) shows the outcome where gene is classified as a cancer gene. The dividing line represents the selected threshold at which the binary decision is optimal (creating a desirable balance between true negatives and true positives). Fig 3 shows how well our model discriminates between prediction classes (cancer gene or non-cancer gene) and shows the selected score (threshold) that could be used to make a binary (true/false) prediction for a gene to be classified as a candidate cancer gene.

**Fig 3.**
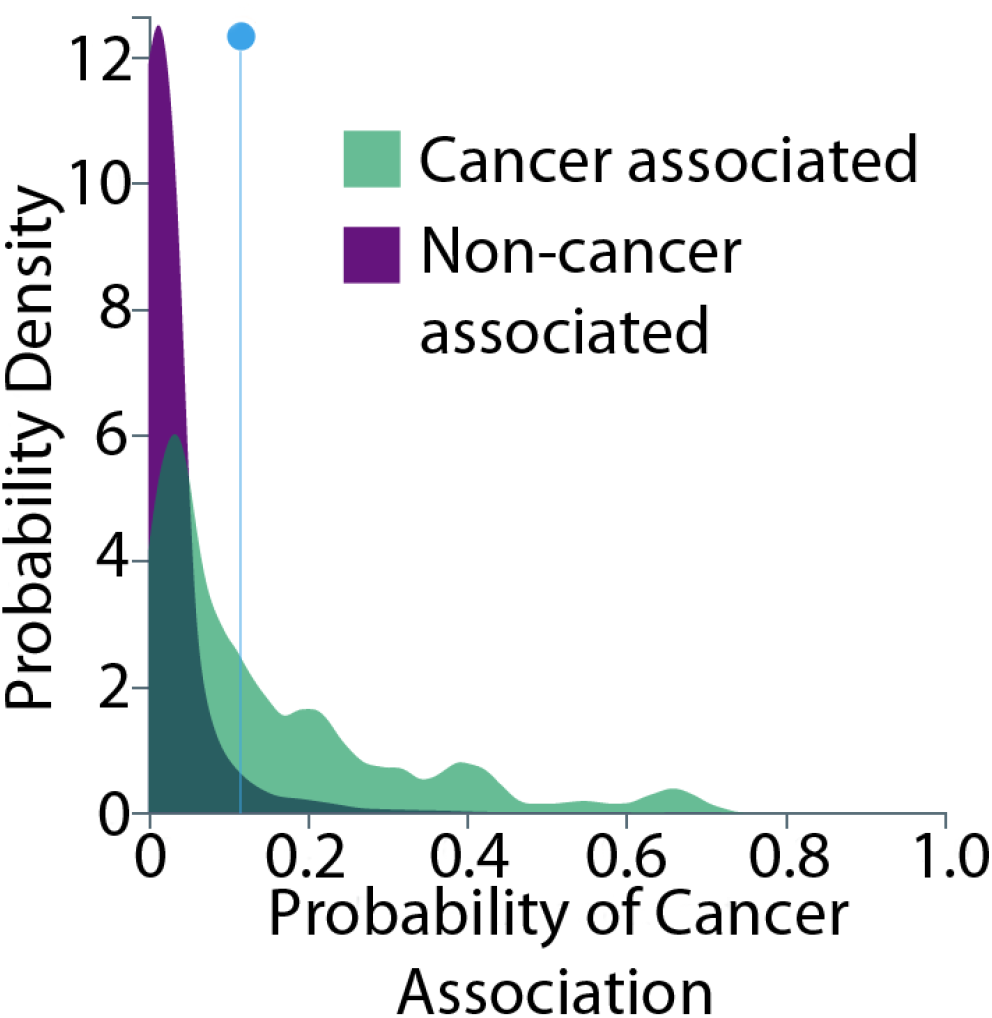
The model’s Prediction Distribution graph showing model performance (how well the model discriminates between cancer and non cancer genes). The colors correspond to the rows of the confusion matrix (Table 4). Every prediction to the left of the dividing line is classified as non-cancer and every prediction to the right of the dividing line is classified as cancer.

The Prediction Distribution graph can be interpreted as follows: purple to the left of the threshold line, is for instances where genes were correctly classified as non-cancer (true negatives). Green to the left of the threshold line is for instances were incorrectly classified as non-cancer (false negatives). Purple to the right of the threshold line, is for instances were incorrectly (according to the current training/validation dataset) classified as cancer genes (false positives and thus potential future cancer genes). Green to the right of the threshold line, is for instances that were correctly classified as cancer genes (true positives). The graph again confirms that the model was able to accurately between cancer and non-cancer genes.

Using the ROC curve produced for our model (Fig 4), we were able to evaluate the accuracy of prediction. The AUC (area under the curve) is a metric for binary classification that considers all possible thresholds and summarizes performance in a single value, with the larger the area under the curve, the more accurate the model. An AUC of 0.5 suggests that predictions based on this model are no better than a random guess. An AUC of 1.0 suggests that predictions based on the model are perfect. (A perfect AUC is highly uncommon and likely flawed indicating that some features that should not be known in advance are being used in model training, thus revealing the outcome). As the area under the curve is of 0.86, we conclude that the model is accurate. The highlighted point on the ROC curve represents the threshold chosen for classification of genes. This is used to transform probabilities scores assigned to each gene into binary classification decision where each gene would be classified into potential cancer genes (true) or not cancer genes (false).

**Fig 4.**
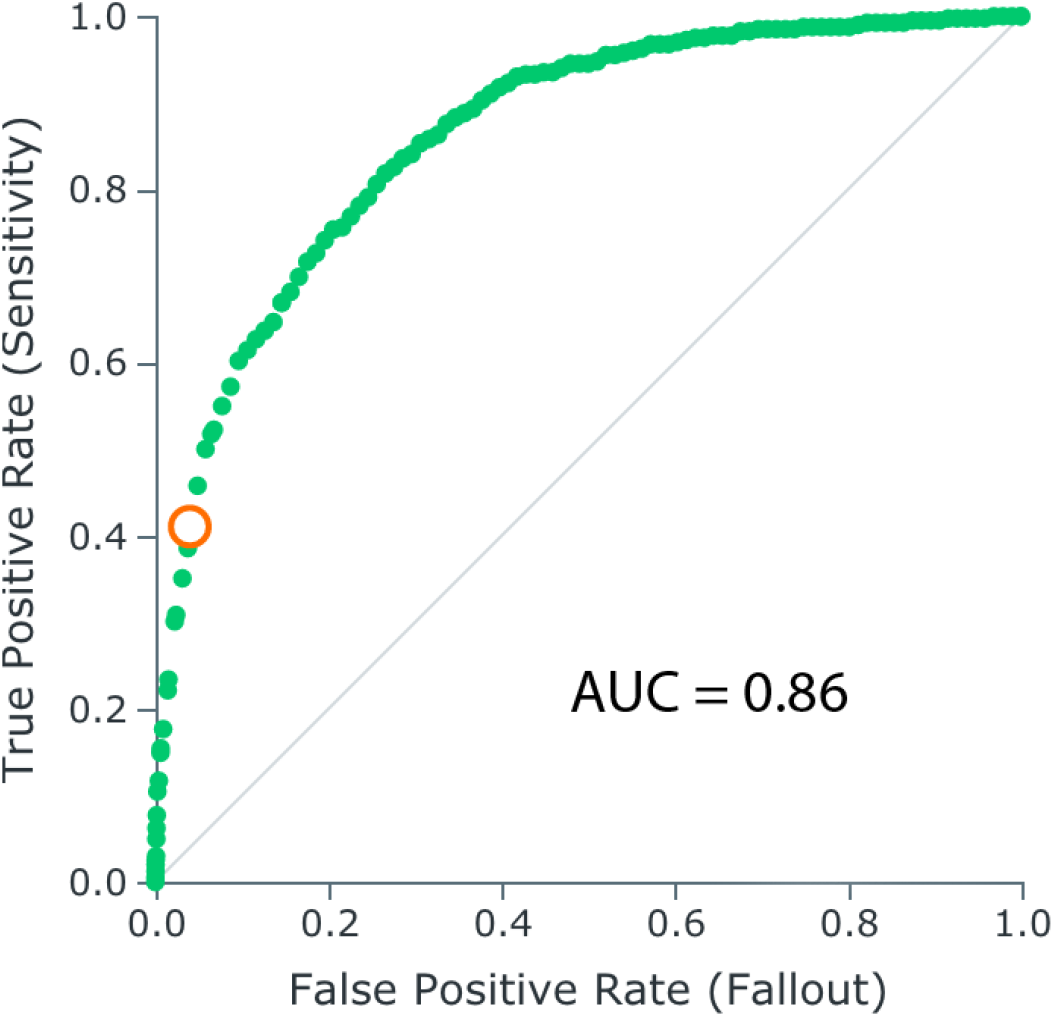
Statistical quality of the model measured using Receiver Operator Characteristic (ROC curve). The orange point shows the threshold for classification of genes.

### Feature Impact

Feature impact shows how useful each feature is for the prediction. It is measured by seeing how much worse a model’s error score would be if the model made predictions after randomly shuffling the values of one field input, while leaving other values unchanged. The platform normalises the scores so that the value of the most important feature is 100%. This helps identify those properties that are particularly important in relation to predicting cancer genes, hence furthering our understanding of the biological aspects underlying the propensity of a gene to be a cancer gene.

‘Closeness’ and ‘Degree’ are the properties with the highest feature impact (Fig 5). Both are protein–protein interaction network properties, indicating a central role of the protein product within the network. We find that both correlate with likelihood of cancer association. Other important properties are the ‘phi’ essentiality score (probability of haploinsufficiency compared to baseline neutral expectation) and Tajima’s D regulatory (measures for genetic variation at intra-species level and for proportion of rare variants). We also note that greater length of a gene or transcript increases the likelihood of a somatic mutation, so increasing the chance of a mutation within that gene, thus increasing the likelihood of it being a cancer gene.

**Fig 5.**
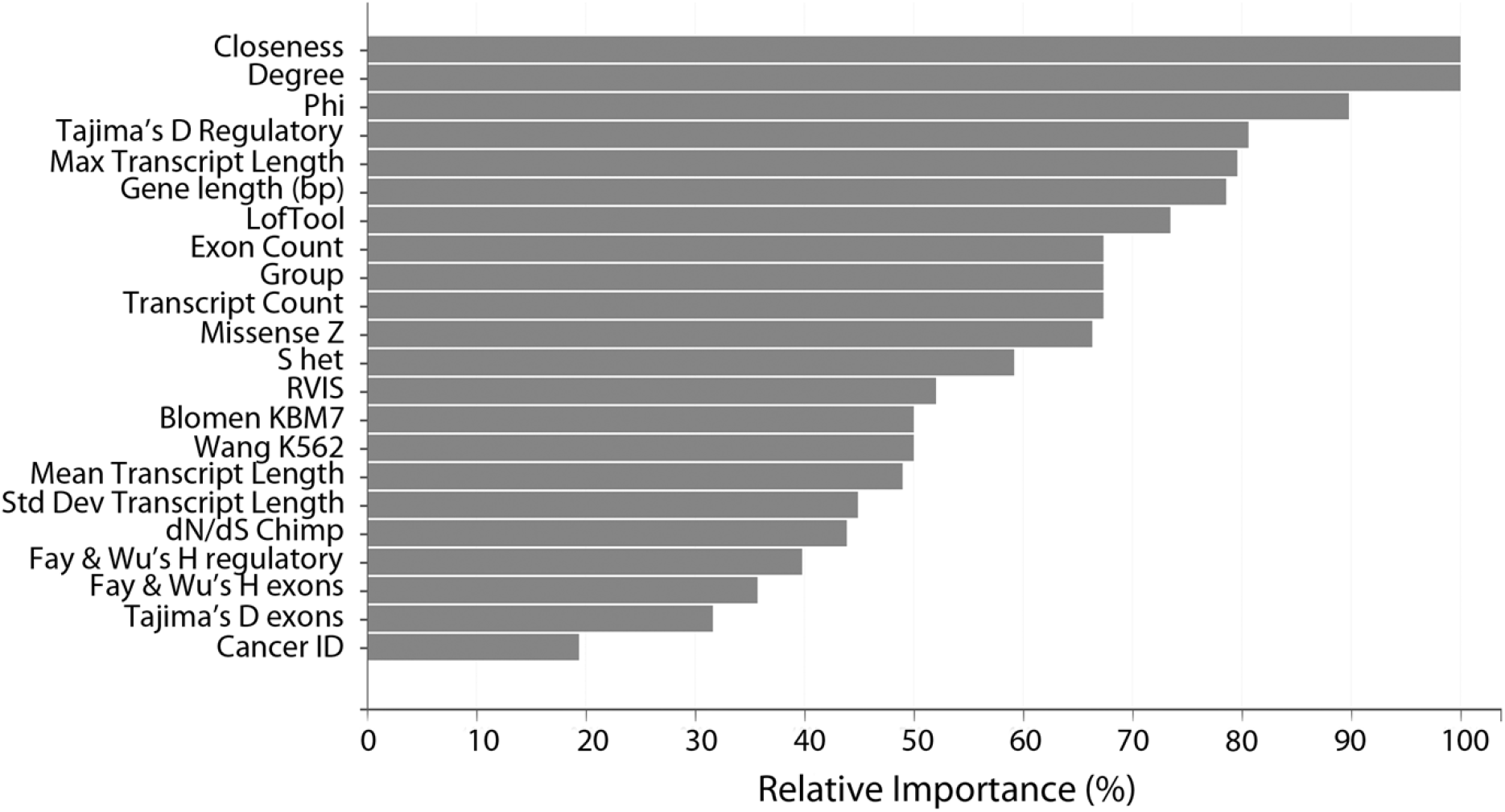
Subset features list selected by the modeling process to be used to make the predictions in our model. Features are shown here ranked by their relative importance (normalised impact of each feature on model’s predictions).

We also retrained our model using a data set that excludes general gene properties, as listed in the ‘Data Sets’ section, and found that a reduction in model’s performance was evident but very small. The model trained on this dataset achieved an AUC of 0.835 and a sensitivity of 55% at a specificity of 89%. This small reduction in the predictability of the models confirms that essentiality and protein-protein interaction network properties are the most important features in predicting cancer genes and that information carried by gene general properties can be in most part be represented elsewhere. For example, longer genes (median transcript length =3737) tend to have the highest number of protein–protein interactions [14], so these properties correlate.

## Discussion

We have demonstrated that gene essentiality is a strong indication that it is associated with cancer. This is the case for both oncogenes and tumour suppressors. Genes classed as essential are often involved in cell, embryo and organism growth. Similarly, proliferation is key for cancer cells. Therefore, the sets of genes that are essential and those that are involved in unregulated growth as seen in cancer tend to overlap.

We used a range of direct methods of essentiality, such as LofTool and Missense Z-score, based on human population sequence data, and Blomen KBM7 and Wang K562 that are based on cell viability data. We also investigated a range of gene and protein properties, such as number of protein-protein interactions and position in the interaction network, expression levels, and various measure of evolutionary selection pressure. All of these gene and protein properties are strongly linked and closely correlate with essentiality, even though they do not measure it directly.

Machine learning allows the identification of the most important features for classification of genes into cancer-related and non-cancer-related groups. Such an approach can provide a reliable framework that first helps in identifying properties predictive of cancer association and can then provide a reliable model that can be used to predict the most likely candidates to be cancer genes. Machine learning methods can produce better performing predictive models than traditional statistical regression methods because they are more flexible and rely on fewer statistical assumptions. A high degree of flexibility in defining the model structure typically results in better model performance. In our application, the only assumption being made is that the model training data is representative of the future scoring data. In our case, this means that current knowledge of cancer genes is applicable to those that will be found in the future. The resulting classifier is accurate (AUC > 0.85) in predicting whether or not a human protein-coding gene is cancer-related.

Informed selection of protein targets is critical when a new cancer drug is developed. Evaluating possible options where increased likelihood of success, such as decreased likelihood of resistance, can be improved with further understanding genetic and evolutionary (selection pressure) characteristics of these genes [15]. To achieve this, widening the pool of cancer gene candidates and increasing understanding of the purifying selection pressure on genes (in this case the degree of essentiality) could help tailor a therapeutic strategy, as they might benefit from being informed with the primary selection pressures that each target gene is subject to.

Our machine-learning model prediction scores provide a good base to prioritise the likelihood of a human protein coding gene to be a cancer gene. Of key importance in our results are those predictions that are false positives – the 1490 genes with high scores that have no published cancer association. Two possible explanations exist: either they represent a failure of the model to correctly classify the data or these gene are in fact cancer related but have not yet been characterised as such. Our success when analysing recently identified cancer genes confirms that these genes are indeed an informed place to look for future cancer targets.

## Methods

### Data Sets

#### Data Source Overview and Appropriateness

A total list of 18,000 human genes was used. Gene properties were obtained by combining data available in previous studies [3, 4] in addition to those available from Ensemble Biomart. Below are the categories and sources of data used:

#### Essentiality scores

We obtained several different essentiality scores calculated for human genes from [4] to use in our dataset. Petrovski’s ‘residual variation intolerance score’ (RVIS) [16] and Rackham’s EvoTol [17] relate the amount of common loss-of-function variation to that of the total gene variation. Other scores are based on the work of Samocha et al. (Missense Z-score)[18], which sets up a baseline expectation of mutation count per gene based on the sequence context, local mutation rate, sequencing depth and, most importantly, sample size. Fadista’s LoFtool [19] combines the neutral mutation rate of Samocha et al. and the evolutionary information in EvoTol. The baseline neutral expectation is compared with the observed counts of loss-of-function variants in the Missense Z-score, in Bartha’s probability of haploinsufficiency (Phi) [20] and in Lek’s probability of loss-of-function intolerance (pLI) [9]. Finally, recent work by Cassa et al. [8] describes a metric (shet) that provides Bayesian estimates of the selection coefficient against heterozygous loss-of-function variation. The various scores were developed or updated using the Exome Aggregation Consortium (ExAC) sample of 60,706 human exomes described in [9]. These scores show high correlations with one another [4]. The key differential characteristics of the scores and all scores’ URLs and values are included in S1 Table.

#### Evolutionary profile and genomic related properties

We used gene properties provided and constructed in [3] including genomic location, protein network parameters and summary statistics of neutrality for human genes.

The Genomic location properties we used in our work were: Chr, Start, End and Strand and dN/dS values which indicates neutrality and selection pressure (multiple species). All were extracted from Ensembl Biomart Genes [21].

Group property divides genes into three different mutually excluding groups: (i) Complex-Mendelian (CM) genes, (ii) Mendelian Non-Complex (MNC) genes, and (iii) Complex Non-Mendelian (CNM) genes.

Data also include measures for genetic variation at intra-species level and measures for proportion of rare variants, such as Tajima’s D exons, Tajima’s D regulatory, Fay and Wu’s H exons and Fay and Wu’s H regulatory [3].

#### Protein network properties

The human protein–protein interaction network (PIN) was reconstructed from the interactions available in the BioGRID database version 3.1.81 [22]. Properties such as degree were computed as the total number of interactions in which a protein is involved, while betweenness and closeness centralities were computed using the NetworkX Python library [23].

#### General gene properties

We enriched the dataset with general gene properties in addition to the properties compiled from the sources above. Properties were directly extracted from Ensembl Biomart Genes [21] were Gene % GC content, Transcript count, Gene Length, while StdDev Transcript length, Average Transcript length, Min Transcript length, Max Transcript length and Exon Count were calculated.

#### Outcome

All genes in our data set were labelled as true or false, indicating whether the gene has been identified as a cancer gene. We did this by identifying if this gene has been added to the Cancer Gene Census in COSMIC [1].

The properties we used to construct our dataset are naturally not inclusive of all possible features. In particular, other studies carried out on mouse have investigated an extended list of essentiality properties, though the subset of features we selected here were shown to be of particular interest (14). Expanding the number of properties used would be an option to explore in the future.

#### Machine Learning platform

We selected the DataRobot [24] machine learning platform (academic version) to conduct our model build. DataRobot simplifies model development by performing a parallel heuristic search for the best model or ensemble of models from a repository of open-source models of different types, based on both the characteristics of the data and the prediction target. This ensures that the selection of our modeling approach is not arbitrary. DataRobot utilises a robust cross-validation and holdout methodology to ensure model performance is sound, reducing the risk of over-fitting.

Machine learning methods can produce more accurate predictive models than traditional statistical methods because they show more flexibility and rely on fewer statistical assumptions. For instance, ordinary least squares regression requires that the Gauss Markov assumptions be supported, to ensure that the model is unbiased and efficient. Traditional statistical regression techniques rely on formal hypothesis testing for variable significance and feature selection (e.g., t-test, p-value, standard error). These statistical tests tend to have assumptions about distribution shape and independence that may not be supported by the data. Machine learning methods, on the other hand, are more flexible in defining the model structure, which typically results in better model performance. Because machine learning includes methods that do not rely on formal hypothesis testing to demonstrate model validity, and because heuristic-style feature selection methods (e.g. stepwise selection) are not used in most machine learning approaches, no such distributional assumptions are required.

#### Machine Learning Algorithm - GBM

The selected model is Gradient Boosting Machines (or Generalized Boosted Models, ‘GBM’). GBM is a cutting-edge algorithm for fitting extremely accurate predictive models [25]. GBMs are a generalisation of Freund and Schapire’s adaboost algorithm (1995) to handle arbitrary loss functions. They are very similar in concept to random forests [25], in that they fit individual decision trees to random re-samples of the input data, where each tree sees a bootstrap sample of the rows of the dataset and N arbitrarily chosen columns where N is a configural parameter of the model. GBMs differ from random forests in a single major aspect: rather than fitting the trees in parallel, the GBM fits each successive tree to the residual errors from all the previous trees combined. This is advantageous, as the model focuses each iteration on the examples that are most difficult to predict (and therefore most useful to get correct). Due to their iterative nature, GBMs are almost guaranteed to over-fit the training data, given enough iterations. The two critical parameters of the algorithm, therefore, are the learning rate (or how fast the model fits the data) and the number of trees the model is allowed to fit. It is critical to cross-validate these two parameters. When done correctly, GBMs are capable of finding the exact point in the training data where over-fitting begins and halt one iteration prior to that. In this manner, GBMs are usually capable of producing the model with the highest possible accuracy without over-fitting.

Our model uses logistic loss and early stopping to determine the best number of trees. Early stopping is a method for determining the number of trees to use for a boosted trees model. The training data is split into a training set and a test set, and at each iteration the model is scored on the test set. If test set performance decreases for 200 iterations (tunable in advanced tuning), the training procedure stops, and the model returns the fit at the best tree seen so far. Note that the early stopping test set will be a 90/10-train/test split within the training data for a given model. The model will therefore internally use 90% of the available training dataset and 10% of the data for early stopping. Since the early stopping test set was used to find the optimal termination point, it cannot be used for training.

#### Model Creation and Validation

To avoid over-fitting, the best practice is to evaluate model performance on out-of-sample data. If the model performs very well on in-sample data, (the training data), but poorly on out-of-sample data, that may be an indication that the model is over-fit.

DataRobot uses standard modeling techniques to validate model performance and ensure that over-fitting does not occur. DataRobot uses a robust model k-fold cross-validation framework to test the out-of-sample stability of a model’s performance. In addition to the cross-validation partitioning, a holdout sample is used to further test out-of-sample model performance and ensure the model is not over-fit. 20% of the training data is set aside as a holdout dataset here. This dataset is used to verify that the final model performs well on data that has not been touched throughout the training process. For further model validation, the remainder of the data is divided into five cross validation partitions. Models that did not perform well at the beginning will not have a cross-validation score. Instead, they will only have a “one-fold” validation.

Because the distribution of the target’s values in a binary classification project may be imbalanced, the validations’ partitions were randomly selected using a stratified sample approach where sub-populations within the data are always represented in each partition to preserve the distribution of the target’s values for each partition.

## Acknowledgments

We thank the leadership team at DataRobot for allowing us to use the platform free of charge for this research.

## Supporting information

**S1 Table. Essentiality scores calculation methods**.

**S2 Table. Essentiality scores for all human protein-coding genes**. This data was used to calculate and show that the cancer genes subset has higher average essentiality scores when compared to the rest of the human genes.

**S3 Table. Machine learning algorithms/models trained on our dataset, with their high-level configuration and performance results**.

**S4 Table. The training dataset used to build the machine learning models** (first tab). Data transformation (preprocessing), such as missing values imputations, and descriptive statistical analysis for all features (numeric and categorical) (second tab).

## Author Contributions

The initial idea for the review article was from A.S. which was closely developed with A.S., S.C.L. and A.J.D. All authors wrote and approved the figures and final wording of the manuscript.

## Ethics declarations and competing interests

Amro Safadi is currently working as a Data Scientist at DataRobot Inc. All the other authors have no conflicts of interest to declare.

## Notes

### Competing Interest Statement

The authors have declared no competing interest.

